# Effect of pancreatic lipase inhibitor and sorbent of lipids on cholesterolaemia and faecal output of fat in rats

**DOI:** 10.1101/2020.08.06.239335

**Authors:** Milan Marounek, Zdeněk Volek, Tomáš Taubner, Dagmar Dušková, Marian Czauderna

## Abstract

Obesity and high cholesterolaemia are major health problems in industrialized countries. The effects of the antiobesity drug orlistat at 0.3 g kg^-1^ and amidated alginate at 40 g kg^-1^ on serum and hepatic cholesterol and the faecal output of fat and sterols were compared in female rats. Rats were fed diets containing cholesterol and palm fat at 10 and 70 g kg^-1^, respectively. Palm fat was provided by coconut meal. Amidated alginate (the octadecylamide of alginic acid) is a sorbent of lipids, and orlistat (tetrahydrolipstatin) is an inhibitor of pancreatic lipase. Both agents significantly increased the faecal loss of fat, orlistat, however, did not significantly decrease serum total cholesterol and its effect on hepatic cholesterol was less pronounced. Amidated alginate at 40 g kg^-1^ significantly decreased serum total cholesterol, LDL cholesterol, hepatic cholesterol, and hepatic lipids, and increased the faecal output of fat and coprostanol (a metabolite of cholesterol). Both orlistat and amidated alginate modified the fatty acid profile in excreted lipids. The concentration of saturated fatty acids decreased and the concentration of unsaturated fatty acids increased. Despite different modes of action, orlistat and amidated alginate were equally efficient in the removing dietary fat from the body. Amidated alginate, however, was more active in the control of serum and hepatic lipid metabolism.

## Introduction

Obesity and high cholesterolaemia are major health problems in industrialized countries. Both obesity and disorders of lipoprotein metabolism are risk factors for chronic diseases (coronary heart disease, cancer, diabetes type 2). Classical treatments for obesity, such as dieting, behavioural modification and exercise, often fail; thus, there is the need for drugs efficient in the treatment of obesity. New drugs are inhibitors of food intake which reduce hunger perception, inhibitors of nutrient absorption, and drugs, which increase energy expenditure [1]. Tetrahydrolipstatin, commercially available as orlistat, is an inhibitor of pancreatic lipase [2]. The alternatives to drugs are soluble fibres (gel-forming polysaccharides) that increase intestinal viscosity and affect the process of digestion and absorption [3]. The cholesterol-lowering activity of soluble fibre has been shown in hypercholesterolemic patients treated with psyllium [4], hydroxypropylmethylcellulose [5], chitosan [6], pectin [7], and alginate [8]. Alginate is a polymer of L-guluronic and D-mannuronic acid, which exists in brown seaweeds [9]. Changing alginate to hydrophobic octadecylamide of alginic acid (amidated alginate) greatly increased its hypocholesterolaemic activity and faecal cholesterol concentration in rats [10]. The aim of the present study was to assess the effect of orlistat and amidated alginate on the faecal output of fat in rats fed a diet supplemented with cholesterol and palm fat. We tested a hypothesis that the effects and of orlistat and amidated alginate are similar. The palm fat, alginate obtained from the supplier, and amidated alginate were characterized in order to obtain information which may be important in the present experiment.

## Experimental

### Animals and diets

Twenty-one female Wistar rats, approximately 6 weeks old, were used. The rats were housed individually in a temperature- and humidity-controlled vivarium (22 ± 1 °C, relative humidity 60 ± 5%). All diets were supplemented with cholesterol at 10 g kg^-1^ and coconut meal at 124 g kg^-1^. The coconut meal containing 56.5% fat was purchased in a healthy food shop.

Table 1 presents the composition of the control and experimental diets. Rat diet ST-1 was supplied by Velaz Ltd. (Lysolaje, Czech Republic). After 4 weeks, the rats were randomly divided into three groups of 7 rats each. Diet no. 2 was supplemented with orlistat at 0.3 g kg^-1^. Diet no. 3 was supplemented with amidated alginate at 40 g kg^-1^ at the expense of cellulose. The orlistat dosing was the average of concentrations used in the experiment of Cruz-Henandez et al. [11]. Diets and water were available *ad libitum*. In the course of this experiment, initial and final body weights were measured. In this experiment, the average initial body weight of the rats was 240 ± 28 g. The study was approved by the Ethics Committee of the Institute of Animal Science and the Central Commission for Animal Welfare of the Ministry of Agriculture of the Czech Republic.

**Table 1.** Composition of control and experimental diets (g kg^-1^)

| Diet | 1<sup>a</sup> | 2 | 3 |
| --- | --- | --- | --- |
| Cholesterol | 10 | 10 | 10 |
| Palm fat<sup>b</sup> | 70 | 70 | 70 |
| Orlistat |  | 0.3 |  |
| Amidated alginate |  |  | 40 |
| Cellulose | 40 | 40 |  |
| Diet ST-1<sup>c</sup> | 866 | 866 | 866 |

<sup>a</sup>Control diet without orlistat or amidated alginate
<sup>b</sup>Coconut meal containing 56.5% fat
<sup>c</sup>The rat diet ingredients were soybean meal, meat and bone meal, cereals, wheat bran, limestone, dicalcium phosphate, salt, and supplements vitamins, trace elements and amino acids. The diet ST-1 contained crude protein, fibre, fat, ash and cholesterol at 224, 45, 33, 43 and $0.29 \text{ g kg}^{-1}$ , respectively.

The experiment duration was 3 weeks, and then, the rats were sacrificed by decapitation after anaesthesia via inhalation of isoflurane (Nicholas Piramal India Ltd., London, U.K.). The rats received 4 g of feed 4 h before they were sacrificed [12].

## Materials and reagents

The sodium salt of alginic acid, low-viscosity product A1112 from brown algae, mannuronic acid sodium, reagent 1-phenyl-3-methyl-5-pyrazolone (PMP), orlistat, a certified reference material (PHR 1445), microcrystalline cellulose and *N*-octadecylamine were purchased from Sigma-Aldrich (Prague). Other chemicals were purchased from P-Lab (Prague, Czech Republic). L-guluronic acid sodium salt was obtained from Carbosynth Ltd., Compton, U.K. The Sylon HTP kit (hexamethyldisilazane-trimethylchlorosilane-pyridine 3:1:9) was purchased from Supelco (Bellefonte, U.S.A.). Isolithocholic acid, norcholic acid, 12-ketolithocholic acid, α-, β-, ω-muricholic acids, and β-sitostanol were purchased from Steraloids Inc. (Newport, U.S.A.). Other sterols were obtained from Sigma-Aldrich (Prague).

### Preparation of amidated alginate

The *N*-octadecylamide of alginic acid was prepared by the reaction of methyl-esterified alginic acid with the *N*-octadecylamine reagent [13]. The reaction with *N*-octadecylamine was performed under heterogeneous conditions. The product was washed with acidified ethanol, petroleum ether, pure ethanol and acetone and finally air dried.

### Analytical methods

#### Titration method for carboxyl group determination

The content of all carboxyl groups *COOH*_t_ (% m/m) in alginic acid and the content of free carboxyl groups *COOH*_f_ (% m/m) of its methyl ester were determined by a titration method [14].

#### Organic elemental analysis and the determination of substitution degree

The contents of C, H and N (% m/m) were determined by organic elemental analysis. The degree of amidation (DA, mol. %) of the final products was calculated according to Taubner et al. [13], based on the carbon and nitrogen content.

#### Molecular weight assay

The reaction with organic dinitro derivatives is based on the ability to reduce dinitrosalicylic acid by reducing polysaccharide groups [15]. Galacturonic acid was used for the calibration.

#### Analysis of the mannuronic to guluronic acid ratio in alginate

Alginate was hydrolysed as described by Lu et al. [16] with some modifications. The derivatization of uronic acids was performed as described by Dai et al. [17]. The same procedure was used to treat the monosaccharide standards and their equimolar mixture. The mannuronate to guluronate ratio was calculated from results of subsequent C18 HPLC-DAD analysis (ratio of peak areas corrected for response factors). An analytical C18 column ZORBAX (Agilent Company) was used.

### Sampling

Samples of mixed blood were drawn from each rat at the time of slaughter. The samples were allowed to stand for 30 min. The sera were separated by centrifugation, stored in a refrigerator, and analysed the subsequent day. After laparotomy, the livers were excised and kept at −40 °C until analysis. During the last 5 days of this experiment, faeces were collected, weighed, pooled, frozen and stored until analysis.

### Analyses of serum, hepatic and faecal lipids

Serum concentrations of cholesterol, triacylglycerols, bilirubin, and activities of aspartate aminotransferase (AST) and alanine aminotransferase (ALT) were determined using commercial kits (BioVendor Ltd., Brno, Czech Republic). The total hepatic and faecal lipids were extracted with chloroform-methanol mixture and measured gravimetrically [18]. Hepatic cholesterol, faecal neutral sterols, and bile acids were determined as described previously [10].

### Other analyses

The fatty acid profile of coconut meal, fat content in the coconut meal, and analyses of the ST-1 diet were performed as described by Skřivan et al. [19]. The fatty acid profile in the excreta was performed according to Marounek et al. [20]. Triacylglycerols in coconut meal were determined as described by Kobayashi et al. [21]. Lipids were dissolved in hexane, insoluble particles removed by centrifugation, and supernatant injected onto the Luna^®^ 3 μm C18(2)100 Å, LC column for separation of hydrophobic compounds (Phenomenex, Torrance, U.S.A.). A HPLC apparatus Shimadzu equipped with evaporative light scattering detector was employed. A gradient program based on two solvents was used: The first solvent consisted of 98.9% hexane, 1% isopropanol and 0.1% acetic acid. The second solvent contained 99.9% isopropanol and 0.1% acetic acid. The method was calibrated with glyceryl tripalmitate dissolved in hexane.

### Statistics

Data analyses were performed by means of the one-way analysis of variance (ANOVA) using the GLM procedure of SAS, version 8.2 (SAS Institute, Cary, NC, U.S.A.). The results are expressed as the mean and standard deviation. Significant differences (*P* < 0.05) were identified using Tukey’s test. The Pearson correlation coefficient was used as a measure of the association between pairs of observations.

## Results

The average final body weights of the rats were 279 ± 16 g. The body weights of the rats did not differ among the treatment groups.

### Characterization of alginate and amidated alginate

Table 2 presents the characteristics of alginate, its methyl ester and its amidated alginate prepared from alginate of low viscosity. The molecular weight of low-viscosity alginate was 58.9 kg mol^-1^. After methylation and amidation, the molecular weight of the final product decreased to 12.3 kg mol^-1^. The average value of the mannuronate/guluronate ratio calculated from the results of three analyses carried out on different days was 2.65 ± 0.05.

**Table 2.**
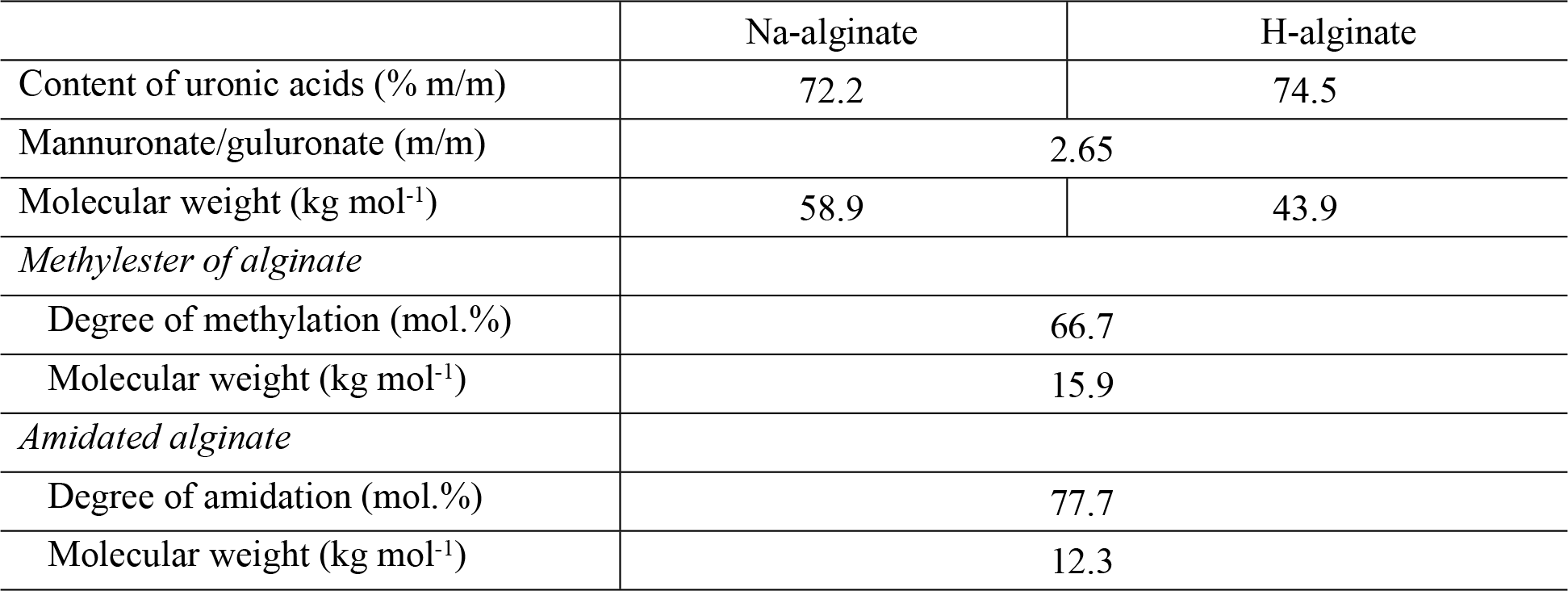
Characterization of alginate, methylester of alginic acid, and amidated alginate

|  | Na-alginate | H-alginate |
| --- | --- | --- |
| Content of uronic acids (% m/m) | 72.2 | 74.5 |
| Mannuronate/guluronate (m/m) | 2.65 |  |
| Molecular weight (kg mol<sup>-1</sup>) | 58.9 | 43.9 |
| *Methylester of alginate* |  |  |
| Degree of methylation (mol.%) | 66.7 |  |
| Molecular weight (kg mol<sup>-1</sup>) | 15.9 |  |
| *Amidated alginate* |  |  |
| Degree of amidation (mol.%) | 77.7 |  |
| Molecular weight (kg mol<sup>-1</sup>) | 12.3 |  |

### Characterization of palm fat (coconut meal)

In coconut meal, the main fatty acid was myristic acid followed by palmitic acid and capric acid. Oleic acid was the main unsaturated fatty acid present in coconut meal (Table 3). Coconut meal contained 56.5% fat as determined by extraction with petroleum ether. HPLC analysis showed that coconut meal contained triacylglycerols expressed as tripalmitin at 623 mg g^-1^.

**Table 3.**
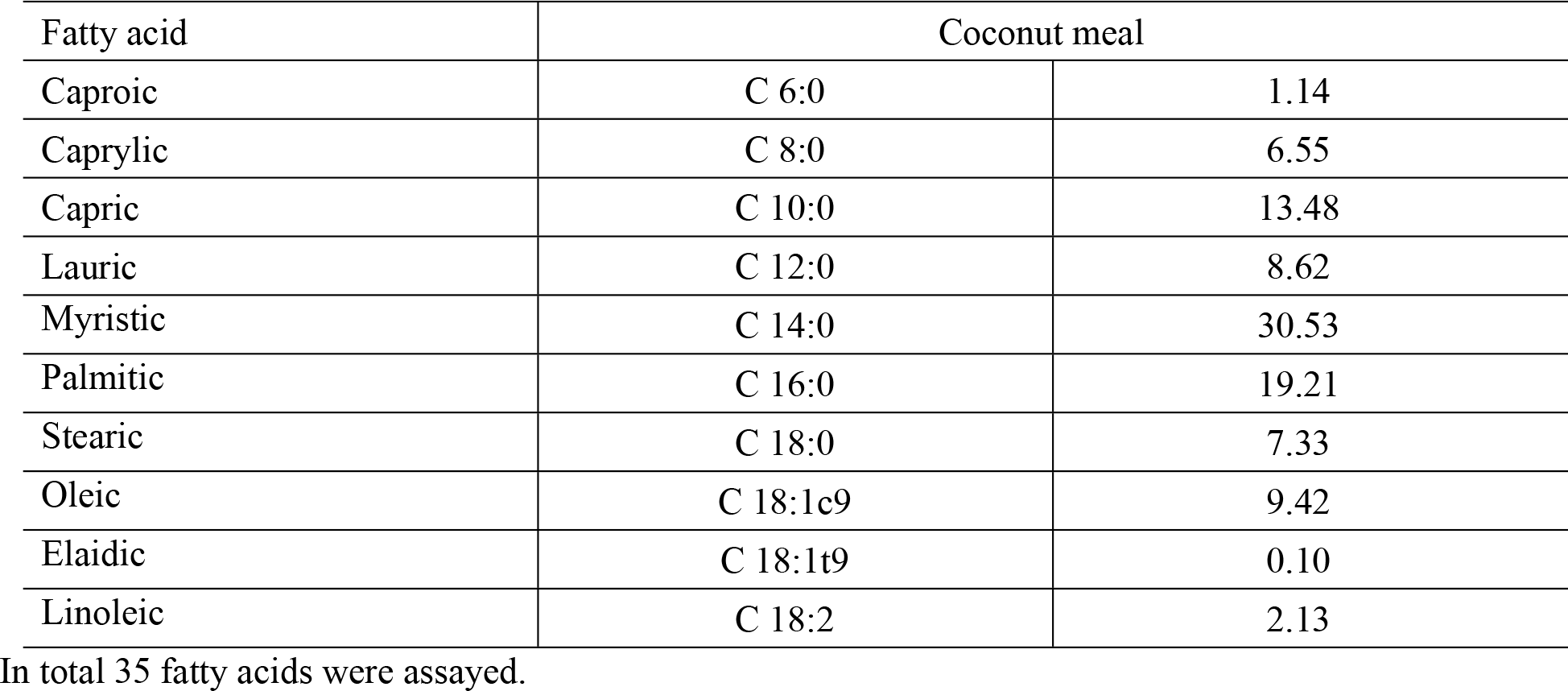
Fatty acid profile of coconut meal (g per 100 g fatty acids determined)

| Fatty acid | Coconut meal |  |
| --- | --- | --- |
| Caproic | C 6:0 | 1.14 |
| Caprylic | C 8:0 | 6.55 |
| Capric | C 10:0 | 13.48 |
| Lauric | C 12:0 | 8.62 |
| Myristic | C 14:0 | 30.53 |
| Palmitic | C 16:0 | 19.21 |
| Stearic | C 18:0 | 7.33 |
| Oleic | C 18:1c9 | 9.42 |
| Elaidic | C 18:1t9 | 0.10 |
| Linoleic | C 18:2 | 2.13 |

In total 35 fatty acids were assayed.

### Effects of orlistat and amidated alginate on serum and hepatic parameters and faecal output of fat and sterols in rats

In rats fed the diet containing 300 mg kg^-1^ orlistat, serum LDL cholesterol, hepatic cholesterol and hepatic lipids significantly decreased; however, total serum cholesterol and serum triacylglycerols were only non-significantly reduced. Supplementation of the rat diet with amidated alginate significantly decreased serum total cholesterol, serum LDL cholesterol, and cholesterol concentration in the liver tissue and the concentration of hepatic lipids. (Table 4).

**Table 4.** Effect of orlistat and amidated alginate (g kg^-1^) on serum and hepatic parameters and faecal output of sterols in rats fed diets 1, 2, 3.

| Diet | 1 | 2 | 3 |
| --- | --- | --- | --- |
| Orlistat |  | 0.3 |  |
| Amidated alginate |  |  | 40 |
| *Serum concentrations* |  |  |  |
| Total cholesterol (μmol mL<sup>-1</sup>) | 3.89 ± 0.29<sup>a</sup> | 2.75 ± 0.90<sup>ab</sup> | 1.75 ± 0.21<sup>b</sup> |
| LDL cholesterol (μmol mL<sup>-1</sup>) | 1.92 ± 0.20<sup>a</sup> | 1.28 ± 0.54<sup>c</sup> | 0.79 ± 0.11<sup>b</sup> |
| Triacylglycerols (μmol mL<sup>-1</sup>) | 2.67 ± 1.53 | 1.71 ± 1.52 | 1.13 ± 0.61 |
| Glucose (μmol mL<sup>-1</sup>) | 8.93 ± 2.52 | 7.26 ± 1.34 | 6.66 ± 0.88 |
| Bilirubin (μmol mL<sup>-1</sup>) | 2.79 ± 0.20 | 2.70 ± 0.32 | 2.71 ± 0.51 |
| *Hepatic parameters* |  |  |  |
| Hepatic cholesterol (μmol g<sup>-1</sup>) | 37.14 ± 8.75<sup>a</sup> | 23.17 ± 3.66<sup>c</sup> | 7.00 ± 0.61<sup>b</sup> |
| Hepatic lipids (mg g<sup>-1</sup>) | 102.4 ± 15.7<sup>a</sup> | 80.9 ± 4.9<sup>c</sup> | 46.3 ± 5.1<sup>b</sup> |
| ALT (nkat mL<sup>-1</sup>) | 1.24 ± 0.32 | 1.53 ± 0.72 | 1.43 ± 0.50 |
| AST (nkat mL<sup>-1</sup>) | 4.22 ± 1.26 | 6.10 ± 3.26 | 3.82 ± 1.18 |
| *Daily faecal output* |  |  |  |
| Fat (g) | 0.42 ± 0.07<sup>a</sup> | 1.00 ± 0.16<sup>b</sup> | 0.97 ± 0.11<sup>b</sup> |
| Cholesterol (μmol) | 481 ± 84 | 581 ± 80 | 569 ± 53 |
| Coprostanol (μmol) | 166 ± 49<sup>a</sup> | 146 ± 52<sup>a</sup> | 245 ± 59<sup>b</sup> |
| Cholesterol + coprostanol (μmol) | 646 ± 108<sup>a</sup> | 726 ± 127<sup>ab</sup> | 814 ± 61<sup>b</sup> |
| Neutral sterols<sup>d</sup> (μmol) | 750 ± 126<sup>a</sup> | 851 ± 140<sup>ab</sup> | 908 ± 67<sup>b</sup> |
| Bile acids (μmol) | 137 ± 22 | 114 ± 30 | 104 ± 28 |
| Total sterols (μmol) | 887 ± 132 | 965 ± 151 | 1012 ± 89 |

Seven rats per group. All diets contained cholesterol and coconut meal
<sup>a-c</sup> Values in the same row with different superscripts differ significantly ( $P < 0.05$ )
<sup>d</sup> Including plant sterols

Serum triacylglycerols correlated significantly with serum cholesterol (*r* = 0.610; *P* = 0.003) and non-significantly correlated with total hepatic lipids (*r* = 0.378, *P* = 0.091). Serum cholesterol correlated significantly with hepatic cholesterol (*r* = 0.804, *P* <10^-4^). The daily faecal output of fat significantly increased in rats fed diets supplemented with amidated alginate and orlistat. Amidated alginate significantly increased the faecal loss of coprostanol and total neutral sterols. The bile acid output, serum concentration of bilirubin, and activity of aminotransferases were not significantly influenced.

Amidated alginate and orlistat modified the fatty acid profiles in excreted lipids. The concentration of saturated fatty acids decreased and the concentration of unsaturated fatty acids increased (Table 5).

**Table 5.** Fatty acid profile^a^ in excreta of rats fed control diet and diets supplemented with orlistat and amidated alginate (g kg^-1^)

| Diet | 1 | 2 | 3 |
| --- | --- | --- | --- |
| Orlistat |  | 0.3 |  |
| Amidated alginate |  |  | 40 |
| Caproic | 0 | 0.40 ± 0.12 | 0 |
| Caprylic | 1.38 ± 0.22<sup>b</sup> | 7.64 ± 0.88<sup>d</sup> | 0.57 ± 0.13<sup>c</sup> |
| Capric | 1.88 ± 0.41<sup>b</sup> | 6.84 ± 0.45<sup>d</sup> | 0.67 ± 0.22<sup>c</sup> |
| Lauric | 29.21 ± 1.67<sup>b</sup> | 18.20 ± 1.83<sup>c</sup> | 16.07 ± 5.80<sup>c</sup> |
| Myristic | 20.75 ± 0.87<sup>b</sup> | 23.31 ± 1.20<sup>b</sup> | 14.56 ± 4.57<sup>c</sup> |
| Palmitic | 16.36 ± 0.56<sup>b</sup> | 14.08 ± 0.34<sup>d</sup> | 22.10 ± 2.53<sup>c</sup> |
| Stearic | 7.13 ± 1.38<sup>b</sup> | 5.88 ± 0.68<sup>b</sup> | 19.78 ± 7.17<sup>c</sup> |
| Oleic | 4.82 ± 0.36<sup>b</sup> | 8.64 ± 0.43<sup>c</sup> | 5.63 ± 0.93<sup>b</sup> |
| Elaidic | 1.16 ± 0.31<sup>b</sup> | 0.70 ± 0.23<sup>b</sup> | 2.72 ± 0.65<sup>c</sup> |
| Linoleic | 7.02 ± 1.43<sup>b</sup> | 9.59 ± 0.59<sup>c</sup> | 6.39 ± 1.66<sup>b</sup> |
| Saturated FA | 85.95 ± 1.89<sup>b</sup> | 80.12 ± 1.35<sup>c</sup> | 77.73 ± 2.68<sup>c</sup> |
| Monounsaturated FA | 6.6 ± 0.65<sup>b</sup> | 9.51 ± 0.55<sup>c</sup> | 8.60 ± 1.07<sup>c</sup> |
| Polyunsaturated FA | 7.37 ± 0.49<sup>b</sup> | 10.23 ± 0.66<sup>bc</sup> | 13.67 ± 1.87<sup>c</sup> |

<sup>a</sup> g per 100 g of fatty acids determined
<sup>b-d</sup> Values in the same row with different superscripts differ significantly ( $P < 0.05$ )

## Discussion

Alginate is a natural polysaccharide, the composition of which depends on the algae species. The molecular weight and the mannuronate to guluronate ratio play a significant role in the physicochemical properties of alginate [16]. During amidated alginate synthesis, acid hydrolysis conditions caused a reduction in the average molecular weight. In the experiment of Sánchez-Machado et al. [22], the mannuronate to guluronate ratio of five seaweed species varied from 2.3 to 4.3. Our findings (2.65 ± 0.05) were thus in this range.

Modified polysaccharides containing long hydrocarbon chains are efficient sorbents of lipids, as shown in experiments with amidated pectin [23], amidated celluloses [24], and amidated alginate [10]. In the present experiment, in rats fed amidated alginate, serum and hepatic cholesterol concentrations decreased, and faecal outputs of fat, cholesterol and coprostanol (cholesterol metabolite) increased. The low concentration of hepatic cholesterol in rats fed experimental diets 2 and 3 may limit the synthesis of bile acids by hepatic 7α-hydroxylase of cholesterol [25].

The aim of this study was to compare effects of orlistat and amidated alginate in rats. Amidated alginate and orlistat increased the faecal loss of fat; however, the mode of action of both agents was different. Amidated alginate is a non-specific sorbent of lipids, and orlistat inhibits triacylglycerol hydrolysis, which is a prerequisite for lipid absorption. It can be concluded that both agents increased the loss of fat from the body, amidated alginate, however, was more efficient in the control of cholesterolaemia and the hepatic cholesterol level. Contrary to amidated alginate, orlistat did not significantly decrease serum total cholesterol and its effect on hepatic cholesterol was less pronounced. Orlistat did not significantly increase the faecal output of cholesterol and its metabolite coprostanol.

Mahmoud and Elnour [26] reported in rats fed a high-fat diet that orlistat at 200 mg/kg significantly decreased serum triacylglycerols, LDL and total cholesterol. Others studies available in the literature on rats mostly focus on the orlistat effects on pancreatic lipase activity, weight gain [26], body fat mass [27], dietary fat attractiveness [28], and treatment of unconjugated hyperbilirubinaemia [29]. Data on the effects of orlistat on the activity of intestinal cholesterol esterase are lacking in the literature. Hydrolysis of cholesterol esters is necessary for cholesterol absorption.

Both amidated alginate and orlistat decreased the concentration of saturated fatty acids in the excreta of rats and increased the concentration of monounsaturated and polyunsaturated fatty acids. This may be related to the fact that unsaturated lipids are more easily incorporated into mixed micelles in the intestine. In our previous study [20] amidated alginate decreased molar percentage of saturated fatty acids in excreta and increased proportion of monounsaturated fatty acids.

The activities of aminotransferases and serum bilirubin did not increase in the rats of the experimental groups; thus, amidated alginate and orlistat did not negatively influence hepatocellular health.

